# Feral Pig Invasion Dynamics in Central Brazil: Human-Driven Introduction, Landscape Drivers, and Ecological Consequences

**DOI:** 10.64898/2026.09.03.749266

**Authors:** Wagner Fischer, Raquel de Faria Godoi Silva, Antonio Conceição Paranhos Filho

## Abstract

Biological invasions by feral swine are among the most important drivers of ecological change worldwide, yet invasion dynamics remain poorly documented in tropical South America. In Brazil, the recent spread of wild boar and wild boar–domestic pig hybrids (*javaporcos*) overlaps with long-established populations of free-ranging feral pigs (*porco-monteiro*) in the Pantanal, creating a unique scenario of dual invasive lineages of *Sus scrofa*. We reconstructed the spatiotemporal invasion process of feral swine (*S. scrofa*) across the state of Mato Grosso do Sul (MS), Brazil, using historical occurrence records, captive-breeding data, landscape variables, detection frequencies, and occupancy estimates from 2007 to 2017. The invasion expanded from 29 municipalities (36.7% of the state) in 2007 to all 79 municipalities by 2017. During this period, at least 247 breeding facilities were identified across 59 municipalities, including numerous informal and clandestine operations involving wild boar and hybrids (*javaporcos*). Municipalities with the largest numbers of breeding facilities and reproductive stocks consistently coincided with areas of highest invasion intensity, indicating that propagule pressure from captive populations strongly influenced feral swine expansion. By 2017, invasive populations occupied approximately 3.2 million hectares (8.85% of the state territory), with the greatest occupancy concentrated in agricultural landscapes dominated by soybean, maize, sorghum, and sugarcane production. Detection frequency and occupied area were positively associated with croplands, exposed soil fields, and water-associated habitats, suggesting that agricultural mosaics linked by riparian corridors facilitate invasion and persistence. Conservative population estimates ranged from 11,850 to 47,400 individuals statewide, although carrying-capacity scenarios suggested populations could exceed 500,000 animals under favorable conditions. Detection patterns involving native peccaries (*Tayassu pecari* and *Pecari tajacu*) indicated differential coexistence responses to invasive swine, with collared peccaries apparently more sensitive to heavily invaded landscapes. Our findings demonstrate that feral swine expansion in central South America was driven by the interaction of anthropogenic propagule pressure, permissive governance, and highly suitable agricultural environments, highlighting the urgent need for coordinated monitoring, regulation of captive breeding, and large-scale adaptive control programs.

## Introduction

The expansion of wild boar (*Sus scrofa*) invasions in Brazil has been recognized since at least 1995, yet management responses remained spatially restricted, intermittent, and insufficient to prevent widespread establishment (Deberdt *et al*. 2005, Simberloff *et al*. 2013, Fischer *et al*. 2017, IBAMA 2017). Only in the past decade has coordinated national-level responsibility for feral pig management emerged (IBAMA 2013), by which time the species had already expanded across diverse regions of the country. This lag between invasion recognition and effective intervention likely facilitated rapid geographic spread, particularly across heterogeneous agricultural landscapes (Pedrosa *et al*. 2015).

In Mato Grosso do Sul, invasion dynamics are further complicated by the coexistence of distinct feral pig lineages of *Sus scrofa*. Long-established populations of free-ranging feral pigs, locally referred to as *porco-monteiro*, originated from feralized domestic stock historically confined to the Pantanal floodplain, whereas recently introduced wild boar and wild boar–domestic pig hybrids (*javaporcos*) have expanded from plateau regions into lowland environments (Keiter *et al*. 2016, Fischer 2018). This convergence raises key questions regarding whether distinct invasion histories and genetic backgrounds influence spatial dynamics, ecological impacts, and interactions with native species (Fischer *et al*. 2017, Sicuro *et al*. 2025).

The invasion success of feral pigs is commonly attributed to broad climatic tolerance, omnivory, behavioral plasticity, and high reproductive rates (Choquenot *et al*. 1996, Merino and Carpinetti 2003, Lowe *et al*. 2004, Long 2005, Parker *et al*. 2013). In Brazil, frequent hybridization between wild and domestic lineages may further enhance adaptive capacity, increasing phenotypic variability and facilitating expansion into novel environments (Deberdt and Scherer 2007, Pedrosa *et al*. 2015, Fischer *et al*. 2017). At the landscape scale, resource availability is expected to be a primary driver of occupancy. Feral pigs require access to water, food, and cover—conditions widely available in the grain-producing regions of Mato Grosso do Sul, where riparian corridors and agricultural mosaics may function as dispersal pathways and resource subsidies (Doutel-Ribas *et al*. 2019).

Human-mediated processes likely play an equally important role. Illegal captive breeding and translocation of animals have been repeatedly implicated in the spread of feral pigs (Deberdt and Scherer 2007, IBAMA 2017), increasing propagule pressure despite longstanding regulatory restrictions (IBAMA 1998). These processes suggest that invasion patterns may reflect not only environmental suitability but also socioeconomic drivers linked to land use and animal management (Wilber *et al*. 2020).

Feral pigs exert widespread ecological and economic impacts, including soil disturbance, vegetation degradation, alteration of water resources, and significant agricultural damage (*e.g.*, Oliver 1993, Cushman *et al*. 2004, Herrero *et al*. 2006, Cuevas *et al*. 2010, Hegel and Marini 2013, Barrios-García *et al*. 2014, Pedrosa *et al*. 2015, Hensel and Hensel 2022, Etges *et al*. 2023). However, their interactions with native fauna remain insufficiently resolved outside the Pantanal, where competition with native peccaries (Tayassuidae) has been most extensively studied (*e.g.*, Desbiez *et al*. 2009, Keuroghlian *et al*. 2009, Galetti *et al*. 2015). In fragmented agroecosystems, emerging evidence suggests potential for niche overlap and negative interactions with native suiforms (Gabor and Hellgren 2000, Salvador 2012, Doutel-Ribas *et al*. 2019), but large-scale assessments are lacking.

Despite rapid expansion, quantitative, spatially explicit estimates of feral pig distribution, occupancy, and population size remain scarce in Brazil. Most existing studies focus on local density or home-range size, limiting inference at broader scales relevant for management (Pedrosa *et al*. 2015).

### Study framework and hypotheses

To address these gaps, we integrate interview-based detection data, spatially explicit occupancy modeling, and environmental and socioeconomic predictors to evaluate the following hypotheses:

- *Spatial expansion hypothesis*: Feral pig distribution reflects an ongoing expansion, with distinct spatial patterns associated with recently introduced wild boar/hybrids (*javaporco*) and long-established free-ranging feral pig populations (*porco-monteiro*).
- *Resource–connectivity hypothesis*: Detection probability, occupancy, and relative abundance increase with landscape features that enhance resource availability and connectivity.
- *Human-mediated spread hypothesis*: Socioeconomic variables linked to land use and animal management increase propagule pressure and shape invasion patterns.
- *Saturation hypothesis*: Population size approaches saturation in long-established regions but remains lower in recently invaded areas.
- *Biotic interaction hypothesis*: Feral pig occurrence is negatively associated with native peccary occurrence, particularly in human-modified landscapes.

We test these hypotheses using a statewide sampling framework combining administrative and spatially explicit units, allowing inference across ecological and management-relevant scales (Figure 1).

**Figure 1.**
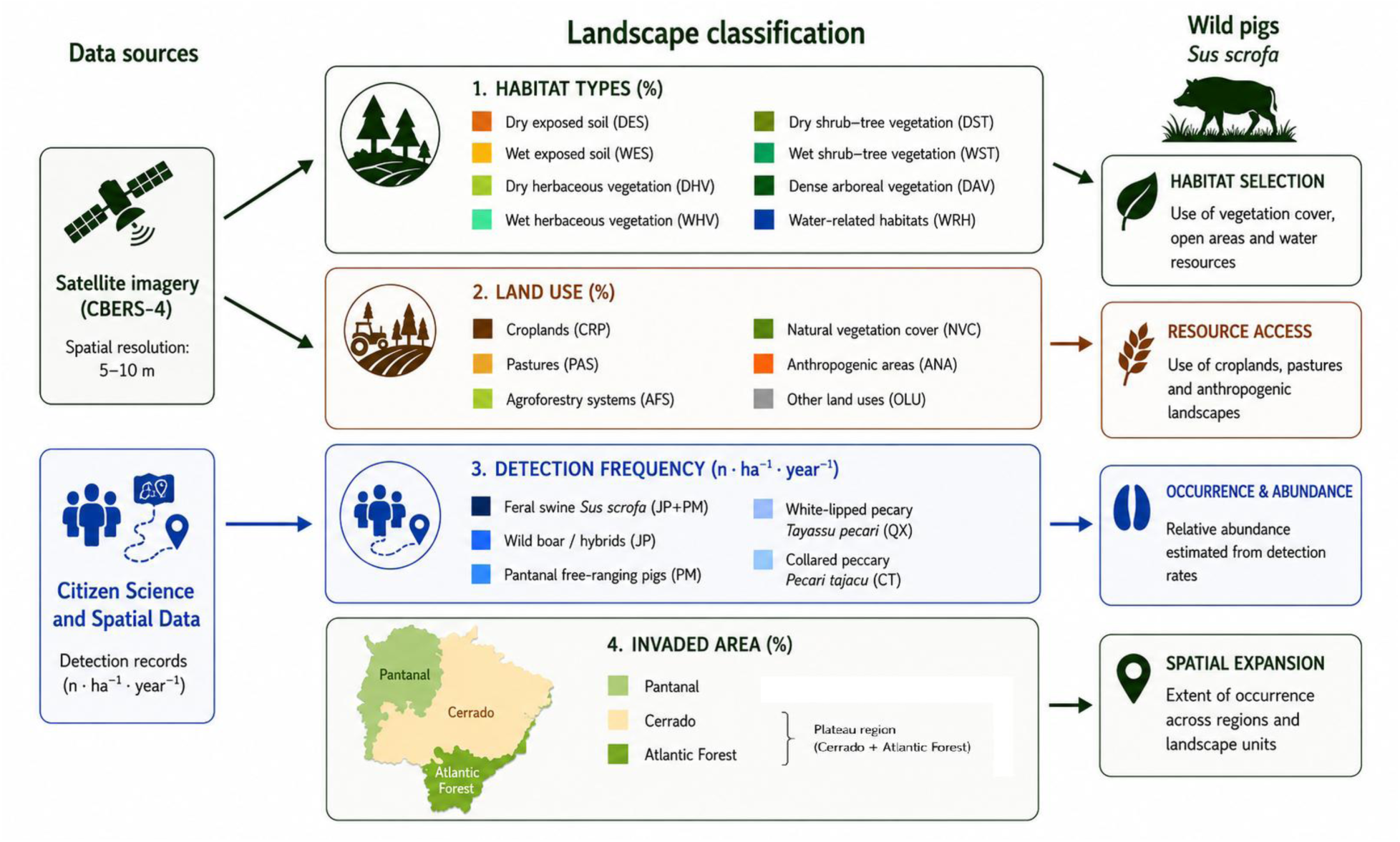
Methodological and conceptual framework of the study. The flowchart illustrates the integration of spatial data (satellite imagery) and citizen science to derive landscape classifications and detection metrics (center). These variables were used to evaluate habitat selection, resource access, and the spatial expansion dynamics of feral swine (*Sus scrofa*) across the Pantanal, Cerrado, and Atlantic Forest biomes (right).

## Methods

### Study area

The study was conducted across the entire state of Mato Grosso do Sul, Brazil (79 municipalities), encompassing approximately 35.7 million hectares of a heterogeneous mosaic of two major landscape domains: the Pantanal floodplain, and the Cerrado and Atlantic Forest plateau ecoregions. This environmental diversity provides a broad gradient for evaluating the effects of landscape structure, resource availability, and human activity on feral pig invasion dynamics. Land use is dominated by agriculture (soybean, maize, sugarcane) and pasturelands, with a dense hydrographic network.

### Sampling design

We adopted a hierarchical sampling framework to link ecological processes with management-relevant spatial units. Municipalities were used as primary sampling units, allowing integration with socioeconomic and policy-relevant data. To account for variation in area, all variables were standardized as proportions. To quantify occupancy and ensure spatial independence, we overlaid a grid of 22,823 hexagonal cells (1,600 ha each). This cell size exceeds reported home-range sizes for feral pigs, minimizing double-counting and allowing each detection to represent independent individuals or groups (*e.g.,* Choquenot *et al*. 1996, Mourão *et al*. 2002, Merino and Carpinetti 2003, Keuling *et al*. 2008, Mitchell *et al*. 2009). This design directly supports estimation of invaded area and spatial extent.

### Data compilation

We compiled data for both invasion processes of *Sus scrofa*, the long-established feral pig populations (*porco-monteiro*=PM) in Pantanal, and the most recent wild boar/hybrids invasion (*javaporco*=JP), from multiple sources:

- Historical records of feral pig occurrence (presence/absence);
- Interviews with local informants (observation/detection of individuals or groups);
- Institutional datasets from: Brazilian Institute of Environment and Renewable Natural Resources (IBAMA), Brazilian Agricultural Research Corporation (EMBRAPA), State Agency for Animal and Plant Health Defense of Mato Grosso do Sul (IAGRO);
- Records of captive breeding facilities of wild boar/hybrids (JP) from 2007 to 2017.

### Environmental and socioeconomic predictors

Landscape structure was quantified using remotely sensed imagery (CBERS-4), from which vegetation (NDVI) and water (NDWI) indices were derived to classify habitat types (INPE 2017). Habitat composition was calculated as proportional area within each sampling unit. Land-use and production variables were obtained from IBGE dataset (2017), including croplands, pastures, agroforestry, natural vegetation, and anthropogenic areas. Major crops were analyzed individually due to their potential influence on resource availability. Additional predictors included livestock abundance, animal production, grain production, and records of wild boar breeding facilities.

### Detection data and interviews

Detection data were obtained through structured interviews conducted statewide (2016– 2017) with individuals regularly exposed to rural environments (see Supporting information: Table S1). The interactive research protocol was approved by the Research Ethics Committee of the Federal University of Mato Grosso do Sul (CEP-UFMS; CAAE 49362315.0.0000.0021; December 2015). Respondents were validated to ensure accurate species identification (White *et al*. 2005), acknowledging that non-detection does not necessarily imply absence (Gu and Swihart 2004). Detection was defined as direct observation, unequivocal signs, or participation in capture/removal events. Detection frequency was classified into ordinal categories and standardized by reported sampling effort (h·year⁻¹). All records were georeferenced and assigned to hexagonal cells, enabling estimation of occupancy as the proportion of invaded cells per municipality (Zeller *et al*. 2011). Respondents also reported occurrences of native peccaries, allowing evaluation of potential biotic interactions.

### Spatiotemporal and population analyses

We reconstructed invasion dynamics using historical snapshots (2007, 2012, 2017) to assess expansion patterns and their association with breeding facilities. Occupancy and detection intensity were mapped at multiple spatial scales. Occurrence data were mapped onto standardized hexagonal sampling units (∼1,600 ha each). Detection frequency (records·ha⁻¹·year⁻¹) and occupancy (percentage of invaded area) were calculated for each municipality. Population size was estimated based on detection rates of observed group sizes and occupied cells (area), and compared with potential population sizes derived from density ranges reported in the literature (0.2–16 individuals·km⁻²), providing bounds for saturation scenarios (Choquenot *et al*. 1996, Bieber and Ruf 2005, Melis *et al*. 2006, Mitchell *et al*. 2009, Barrios-García and Ballari 2012).

### Statistical analyses

We used generalized linear models (GLMs) to evaluate relationships between predictor variables and response variables (detection frequency and occupancy). Candidate models were developed a priori based on ecological hypotheses and compared using Akaike’s Information Criterion corrected for small sample sizes (AICc). Competing models were evaluated according to explanatory power (R²), parameter significance, and ecological interpretability (Pepin *et al*. 2022). Analyses assumed that validated respondents provided reliable detection data and that variation in detection reflected ecological processes after accounting for sampling effort.

## Results

### Spatiotemporal dynamics of invasion

The reconstruction of the invasion process revealed a rapid expansion of invasive feral pig populations across Mato Grosso do Sul between 2007 and 2017 (Figure 2). In 2007, feral pigs were recorded in 29 municipalities (36.7% of the state), with clear spatial segregation between long-established Pantanal feral pigs (PM) and recently established wild boar/wild pig hybrid populations (JP). By 2012, the invasion had expanded to 53 municipalities (67.1%), and by 2017 invasive populations were recorded in all 79 municipalities of the state, indicating complete statewide occupation (Figure 2).

**Figure 2.**
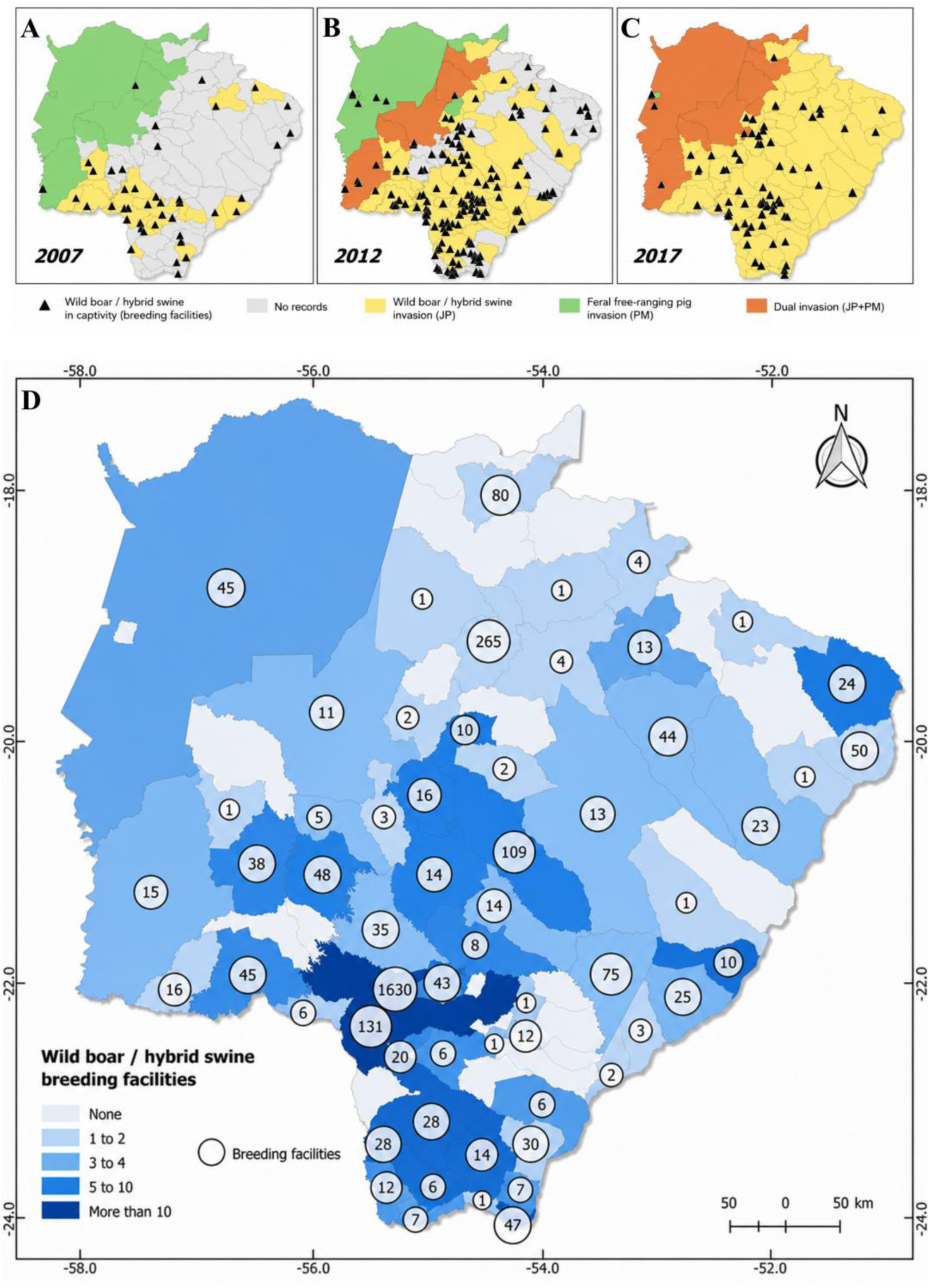
Spatiotemporal progression of feral swine (*Sus scrofa*) invasion in Mato Grosso do Sul, Brazil, and distribution of captive breeding facilities. (A–C) Temporal progression of invasion from 2007 to 2017, showing the transition from isolated foci to statewide occupation. Black triangles indicate wild boar/hybrid swine captive breeding facilities in each period. (D) Consolidated distribution and density of wild boar/hybrid swine breeding facilities per municipality, highlighting the concentration of propagule sources in southern and central agroecosystems. Blue gradient indicates the number of facilities per municipality; proportional circles indicate total facility counts.

The temporal expansion of feral populations closely paralleled the distribution of captive breeding facilities (Figure 2). The number of JP breeding operations increased from 43 facilities in 2007 to 191 in 2012, followed by a reduction to 91 facilities by 2017. This spatiotemporal pattern indicates that each wave of invasion into previously unoccupied areas was consistently preceded by the prior existence of breeding operations. Accordingly, municipalities historically concentrating the largest numbers of breeding facilities and reproductive stock—including Dourados, Ponta Porã, São Gabriel do Oeste, and Bela Vista—later became important invasion hotspots.

### Distribution and occupancy

Feral pigs currently occupy approximately 3.2 million hectares in Mato Grosso do Sul, corresponding to 8.85% of the state territory (Figure 3). Occupancy was spatially heterogeneous, with the highest detection frequencies concentrated primarily in southern and central agricultural regions associated with intensive crop production. Municipalities such as Fátima do Sul, Glória de Dourados, Maracaju, Dourados, Laguna Carapã, and São Gabriel do Oeste exhibited the highest detection frequencies and largest occupied areas (Figure 3).

**Figure 3.**
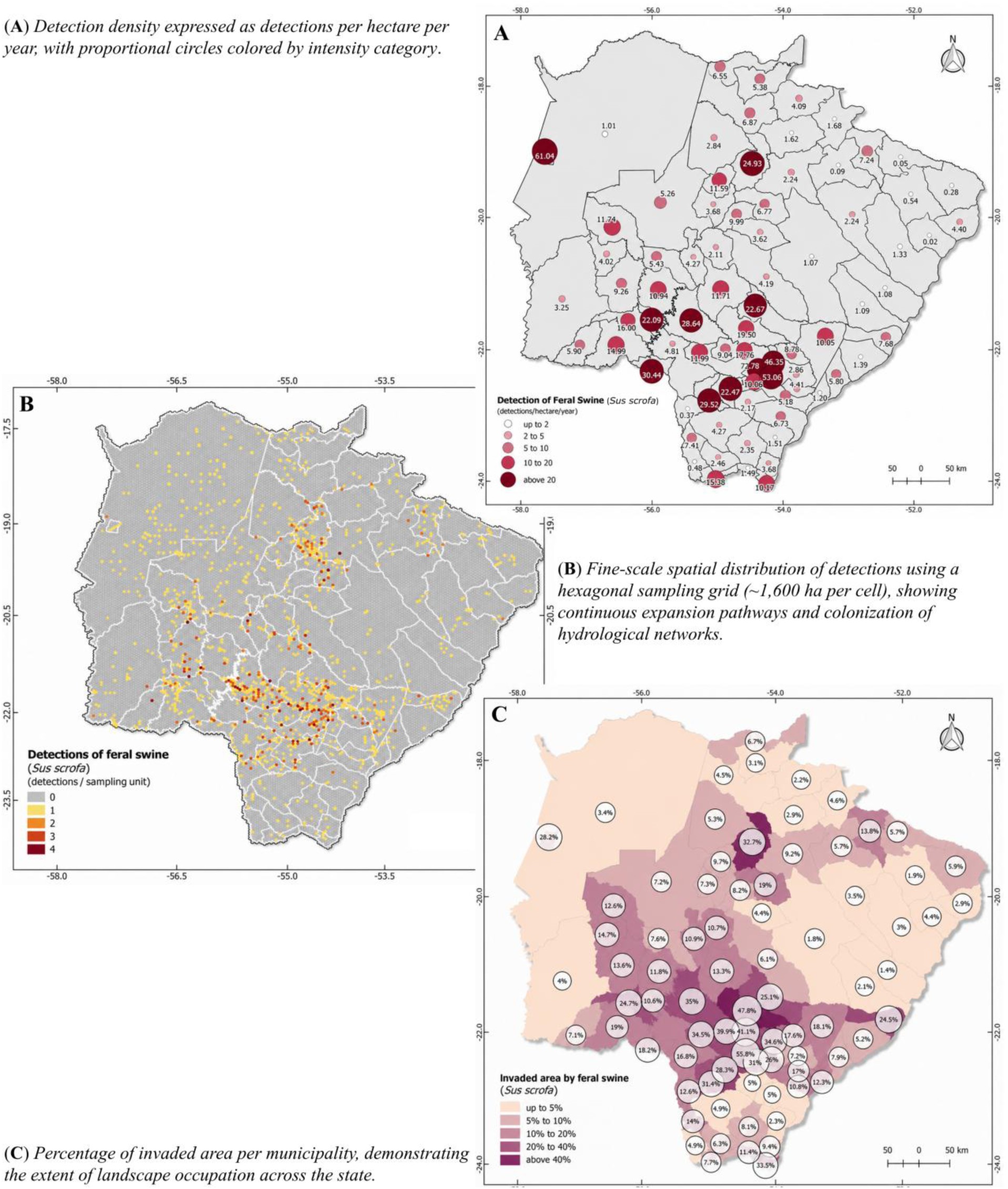
Detection metrics and landscape occupation by feral swine (*Sus scrofa*) in Mato Grosso do Sul, Brazil. Each panel uses a distinct color palette to differentiate the three metrics.

At the landscape scale, plateau agroecosystems exhibited substantially greater occupation than the Pantanal floodplain. Invasion hotspots were strongly associated with regions dominated by soybean, maize, sorghum, and sugarcane cultivation, whereas lower occupation frequencies were generally observed in more isolated municipalities and portions of the Pantanal. Spatial patterns also followed major hydrological and drainage corridors, suggesting that riparian systems contribute to landscape connectivity and dispersal of invasive populations.

### Population Estimates

Population estimates derived from occupied areas indicated a conservative statewide population of approximately 29,600 feral pigs, ranging from 11,850 to 47,400 individuals among municipalities (Figures 4-5). The largest estimated populations occurred primarily in highly productive agricultural regions and portions of the Pantanal, particularly in municipalities such as Corumbá, Rio Brilhante, Maracaju, Dourados, and São Gabriel do Oeste.

**Figure 4.**
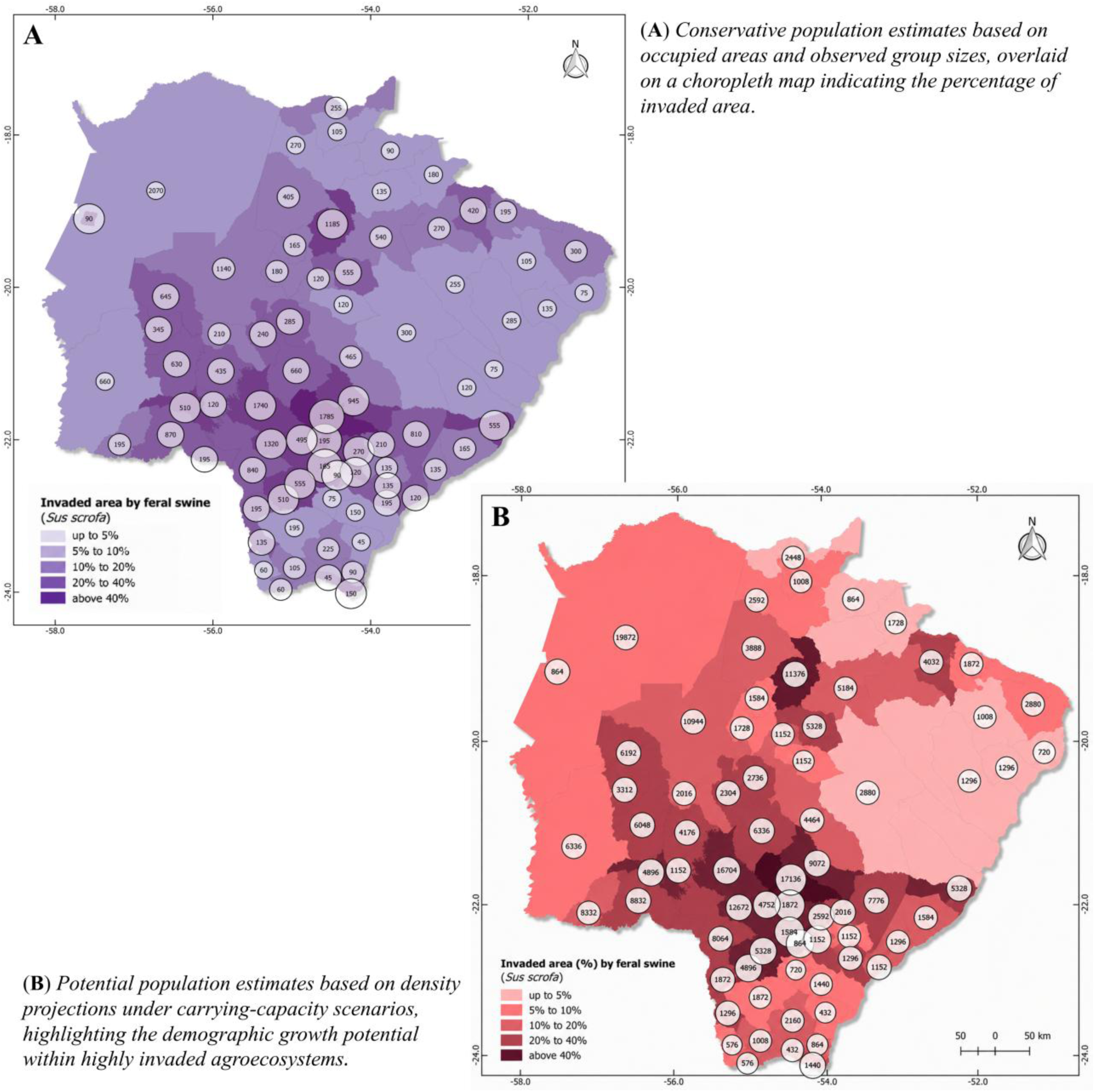
Population estimates of feral swine (*Sus scrofa*) per municipality in Mato Grosso do Sul. Both panels share the same choropleth base (invaded area %) but use distinct color schemes (purple for conservative, red for potential) to differentiate estimation approaches.

**Figure 5.**
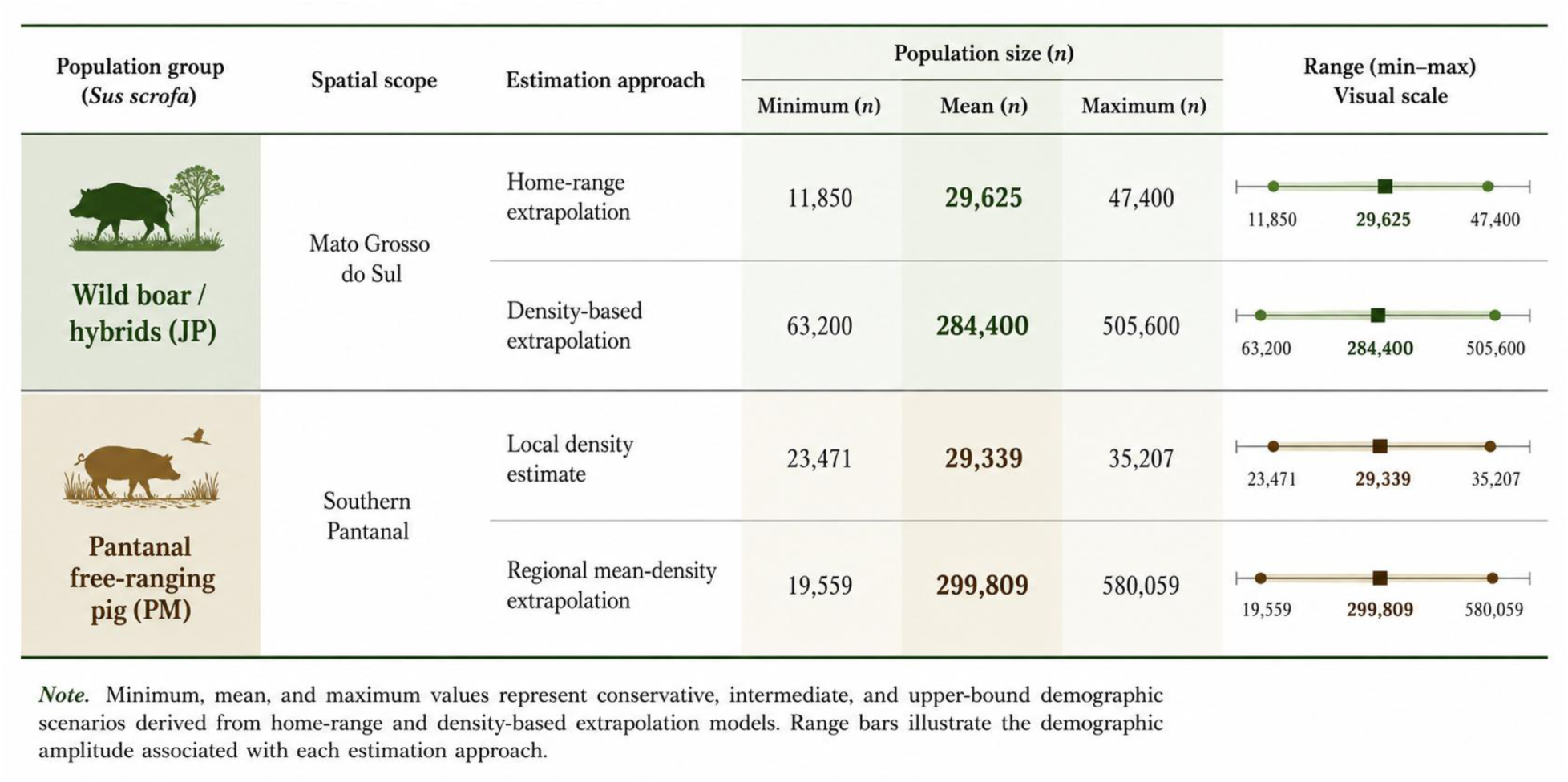
Graphical representation of conservative and density-based population estimates for two *Sus scrofa* lineages in Mato Grosso do Sul: wild boar and hybrids (*javaporco*, JP) at the statewide scale, and Pantanal free-ranging pigs (*porco-monteiro*, PM) in the Southern Pantanal. Minimum, mean, and maximum values represent conservative, intermediate, and upper-bound demographic scenarios derived from home-range and density-based extrapolation models. Range bars illustrate the demographic amplitude associated with each estimation approach.

Density-based projections indicated substantially greater invasion potential (Figure 4). Under average density scenarios, the invaded landscape could support approximately 284,000 individuals, whereas high-density scenarios projected populations exceeding 500,000 animals statewide (Figure 5). These estimates suggest that much of the invaded landscape has not yet reached ecological carrying capacity, especially within intensive agricultural systems characterized by abundant food resources and permanent water availability.

Population estimates for Pantanal free-ranging pigs (PM) revealed a similarly substantial demographic reservoir within the floodplain. Local density estimates for the Southern Pantanal indicated a PM population of approximately 29,300 individuals (range: 23,471–35,207), while regional mean-density extrapolations projected mean populations of approximately 299,800 individuals, with upper-bound estimates exceeding 580,000 animals (Figure 5). Under conservative scenarios, PM populations in the Pantanal alone were thus comparable to statewide JP populations (∼29,600 individuals), whereas under density-based scenarios, PM estimates substantially exceeded JP projections, underscoring the demographic significance of this long-established *Sus scrofa* lineage in the Pantanal wetland.

### Environmental drivers of invasion

Landscape analyses demonstrated strong associations between feral pig occurrence and agricultural land use (Figures 6–7). 26% of the variation in feral swine detection rates (n·ha⁻¹·year⁻¹) was explained by variation in the proportional cover (% area) of three habitat types: dry exposed soil (DES), wet herbaceous vegetation (WHV), and wet shrub–tree vegetation (WST) (F₃,₇₅ = 8.9; r² = 0.26; p < 0.001; Detection = 0.40 DES − 0.29 WHV + 0.49 WST; standardized coefficients). In addition, 41% of the variation in the proportion of invaded area (occupancy) by feral swine among municipalities was explained by variation in the cover of DES and water-related habitats (WRH) (F₂,₇₆ = 26.7; r² = 0.41; p < 0.001; Invaded Area = 0.67 DES + 0.16 WRH; standardized coefficients).

**Figure 6.**
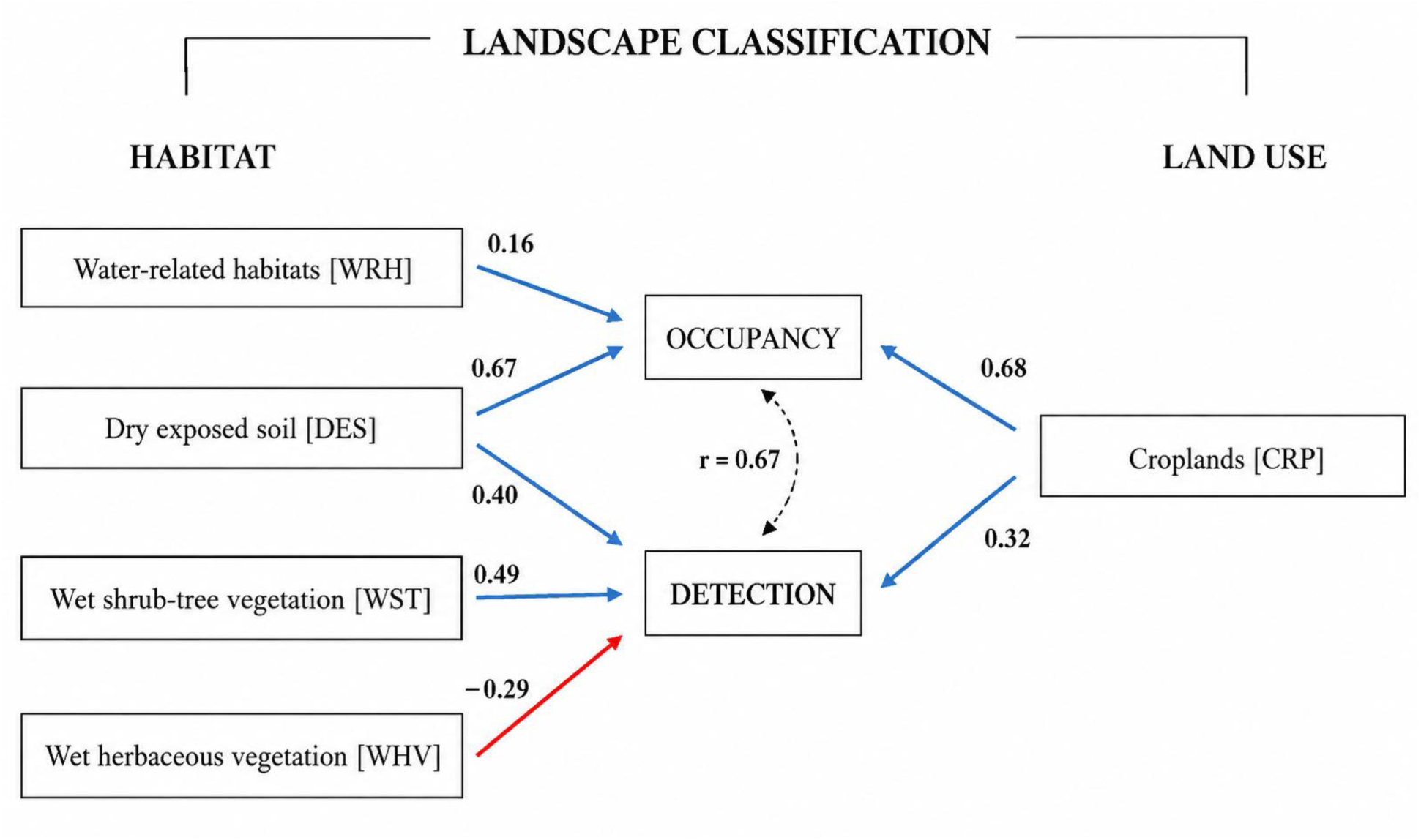
Path analysis diagram illustrating the effects of landscape classification (habitat types and land use) on the occupancy and detection probabilities of feral swine. Blue arrows indicate positive associations, the red arrow indicates a negative association, and numerical values represent standardized path coefficients.

**Figure 7.**
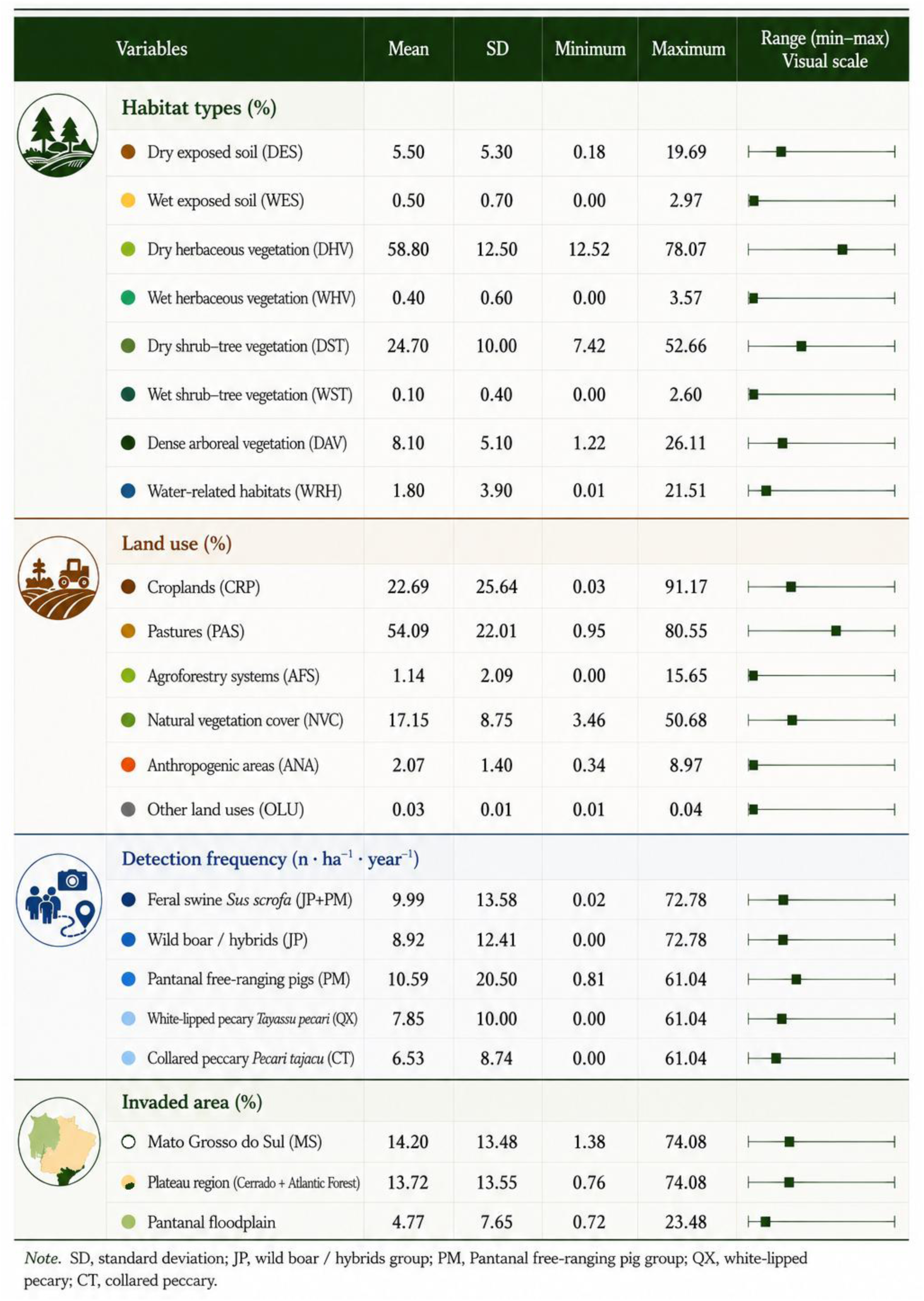
Visual summary of descriptive statistics for landscape composition, land-use categories, and detection frequency of suiforms. The graphical scale (right) illustrates the range (minimum-maximum) and mean values across sampling units in Mato Grosso do Sul and the Pantanal floodplain.

Based on land-use classification, 10% of the variation in feral swine detection frequency among municipalities was explained by variation in cropland cover (CRP) (F₁,₇₇ = 8.6; r² = 0.10; p < 0.01; Detection = 0.32 CRP; standardized coefficient). Most notably, cropland cover explained 46% of the variation in the proportion of feral swine invaded area among municipalities (F₁,₇₇ = 64.3; r² = 0.46; p < 0.0001; Invaded Area = 0.68 CRP; standardized coefficient). In both models, for detection frequency and invaded area (occupancy), the variables natural vegetation cover (NVC), agroforestry systems (AFS), anthropogenic areas (ANA), and other land uses (OLU; residual category) were excluded during model selection procedures. Pasturelands (PAS) were excluded prior to analysis because they were strongly correlated with CRP (r = −0.93; p < 0.001), which showed a positive effect. Across municipalities, feral swine detection frequency (n·ha⁻¹·year⁻¹) was positively correlated with the proportion of invaded area (occupancy) (r = 0.67; p < 0.001), as expected and consistent with the fact that both variables responded similarly to variation in DES and CRP (Figures 6–7).

The distribution (expressed as detection frequency) of feral swine populations (wild boar/hybrid swine and free-ranging feral pigs) and their occupancy (expressed as invaded area) in Mato Grosso do Sul was tested against landscape-environmental variables (Figure 7). The models that best explained feral swine distribution and occupancy across the state responded consistently to dry exposed soil (DES) habitats and cropland land use (CRP), which effectively represent the same landscape type at different temporal stages (Figure 6).

Detection frequencies were positively associated with freshly tilled soils, moist shrub-tree habitats, and drainage networks, whereas moist herbaceous habitats showed negative associations with invasion intensity. These results indicate that agricultural mosaics provide highly favorable environmental conditions for feral pigs by combining predictable food availability, shelter, and permanent access to water.

### Interactions with native suiform species

The four suiform species examined in this study are illustrated in Figure 8. Detection frequencies of invasive feral pigs and native tayassuids revealed asymmetric occurrence patterns among species (Figure 9). White-lipped peccaries (*Tayassu pecari*) were more frequently detected than collared peccaries (*Pecari tajacu*) throughout the state, whereas collared peccaries exhibited the highest number of local absences. Invasive pigs were recorded in all municipalities sampled, including four municipalities where native tayassuids were absent.

**Figure 8.**
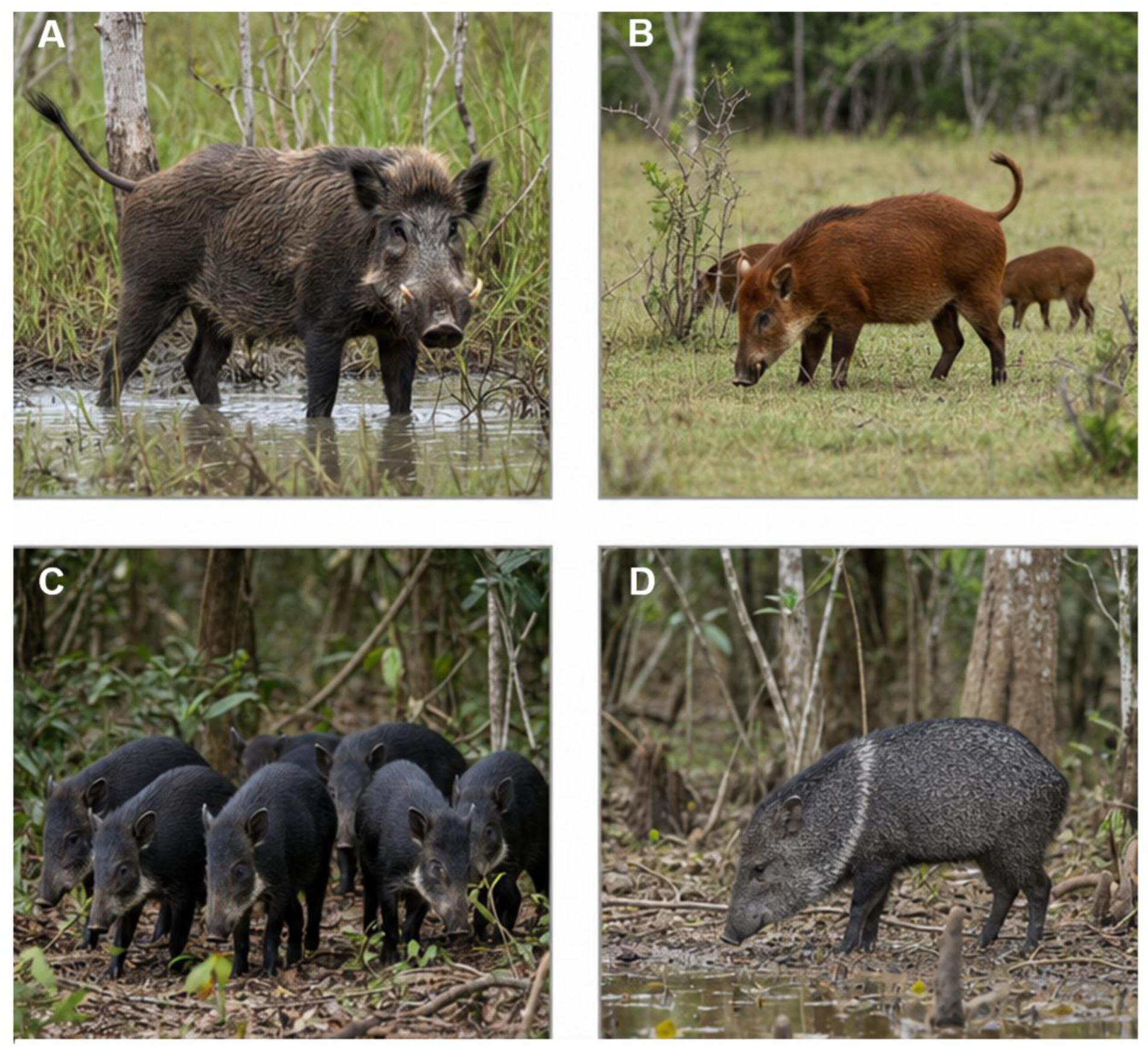
Representative images of the two invasive *Sus scrofa* lineages and the two native peccary species in Mato Grosso do Sul, Brazil. (A) Wild boar and wild boar–domestic hybrid (*javaporco*, JP), recently introduced lineage expanding from plateau regions; (B) Free-ranging feral pig (*porco-monteiro*, PM), long-established Pantanal population derived from feralized domestic stock; (C) White-lipped peccary (*Tayassu pecari*); (D) Collared peccary (*Pecari tajacu*). Composite plate assembled by the authors from reference photographs, with AI-assisted image editing used for compositional refinement and morphological standardization. Credits: Wagner Fischer.

**Figure 9.**
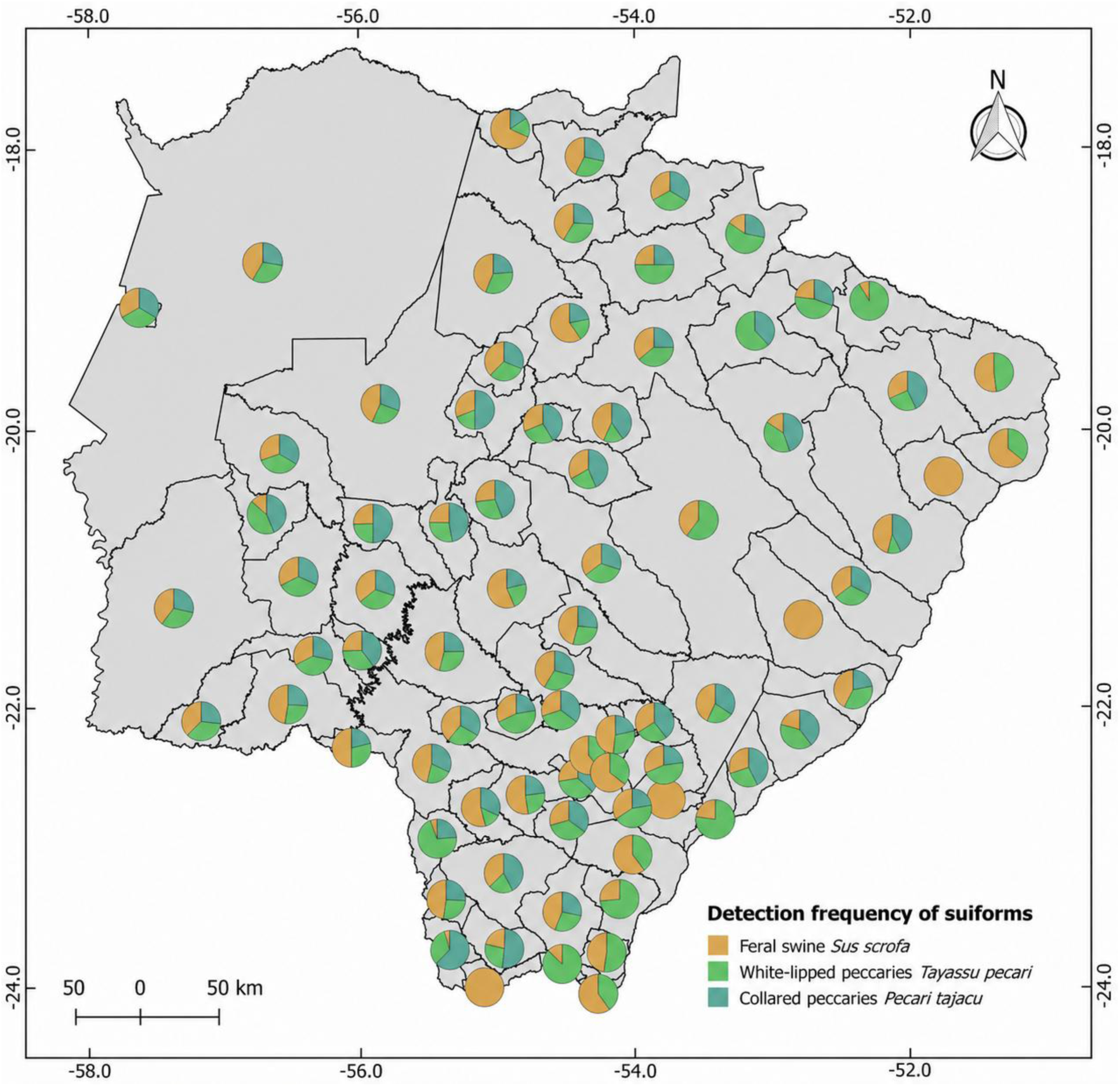
Spatial overlap and detection frequency of suiform species across the study area. Proportional pie charts represent the relative detection rates of invasive feral swine (*Sus scrofa*) compared to native white-lipped peccaries (*Tayassu pecari*) and collared peccaries (*Pecari tajacu*).

A consistent pattern emerged in invaded areas: collared peccaries were never detected in the absence of white-lipped peccaries, whereas white-lipped peccaries frequently persisted even under high feral pig detection frequencies. Although occupancy analyses detected no significant statistical relationship between invasive pigs and native tayassuid occurrence, field observations and detection patterns suggested potential ecological interactions involving habitat overlap, behavioral interference, and possible competitive effects. These patterns were particularly evident within fragmented agricultural landscapes where invasive pigs and native peccaries frequently used the same crop fields, riparian forests, and feeding areas.

## Discussion

### Spatiotemporal invasion dynamics of Sus scrofa: breeders as primary propagule sources

The invasion of *Sus scrofa* in Mato Grosso do Sul followed a multistage process involving two invasive lineages: the long-established Pantanal free-ranging pig (*porco-monteiro*, PM) and the more recent expansion of wild boars and hybrids (*javaporcos*, JP). While PM populations originated from domestic pigs that became feral in the Pantanal nearly two centuries ago (Fischer *et al*. 2017, Pedrosa *et al*. 2021), the JP invasion expanded rapidly across plateau regions during recent decades (Deberdt and Scherer 2007, Pedrosa *et al*. 2015, IBAMA 2017, Fischer 2018).

The temporal reconstruction revealed a strong association between captive breeding and the establishment of free-ranging populations. In 2007, feral pigs occurred in only 29 municipalities, whereas 43 breeding facilities were already distributed across the state, including municipalities without records of feral populations. By 2012, the invasion had expanded to 53 municipalities alongside a marked increase in breeding facilities, strongly suggesting that propagule pressure generated by breeding, trade, escapes, and intentional releases was a major driver of expansion (Deberdt and Scherer 2007, IBAMA 2017). By 2017, feral pigs had been recorded in all 79 municipalities, indicating complete statewide occupation and increasing spatial overlap between PM and JP lineages, particularly in the Pantanal.

This trajectory follows the canonical stages of biological invasions — introduction, establishment, expansion, and saturation (Lockwood *et al*. 2007) — while also highlighting continuous anthropogenic reinforcement throughout the process (Fischer *et al*. 2017) with potential for cross-border dynamics given the state’s international borders (Jaksic *et al*. 2002). Weak enforcement of breeding restrictions, clandestine trade, and hybridization practices likely accelerated invasion dynamics (Deberdt and Scherer 2007, IBAMA 2017, Fischer 2018). The increasing overlap between historical Pantanal feral pigs and recently introduced wild boars may represent a new phase of ecological and genetic admixture with uncertain long-term consequences for the Pantanal ecosystem (Sicuro and Oliveira 2002, Mallet 2005, Fischer *et al*. 2017, Pedrosa *et al*. 2021, Sicuro *et al*. 2025).

### Distribution and landscape-level dominance of feral pigs

Feral pigs are now present in all municipalities of Mato Grosso do Sul, although invasion intensity remains spatially heterogeneous. The highest detection frequencies occurred primarily in southern and central agricultural regions, including Fátima do Sul, Glória de Dourados, Maracaju, São Gabriel do Oeste, and Dourados, whereas lower frequencies persisted in more isolated municipalities and parts of the Pantanal.

Despite statewide occurrence across all municipalities, the effectively invaded area currently corresponds to approximately 3.2 million hectares (8.85% of the state territory), concentrated primarily in agricultural landscapes outside the Pantanal floodplain. Agricultural mosaics dominated by soybean, maize, sorghum, and sugarcane appear to provide highly favorable conditions for establishment and expansion through abundant food resources, habitat heterogeneity, and permanent access to water (Choquenot *et al*. 1996, Choquenot *et al*. 1997, Choquenot 1998, Doutel-Ribas *et al*. 2019). Landscape models reinforced this interpretation, with croplands, tilled soils, and drainage networks explaining much of the observed variation in occupancy and detection frequency.

The spatial distribution of invasion hotspots closely mirrored the historical concentration of captive breeding facilities. Municipalities that historically concentrated breeders and reproductive stock—such as Dourados, Ponta Porã, São Gabriel do Oeste, and Bela Vista— currently exhibit some of the largest occupied areas and highest detection frequencies. Together, these patterns indicate that invasion dynamics in Mato Grosso do Sul are strongly structured by the interaction between anthropogenic propagule pressure, agricultural expansion, and landscape connectivity.

### Population size and environmental drivers of the invasion

Croplands (CRP) in Mato Grosso do Sul are dominated by soybean (48.6%), maize (35.4%), and sugarcane (14.2%) plantations, all highly attractive to invasive feral swine, particularly maize, for which crop damage and economic losses are recurrent and well documented (*e.g.*, Amici *et al*. 2012, Batista 2015, Pedrosa *et al*. 2015, IBAMA 2017, Doutel-Ribas *et al*. 2019). Together with pasturelands, these crops dominate land use across the state (Figure 7).

Dry exposed soil (DES) represents temporary agricultural landscapes during fallow periods, immediately after harvest, or during soil preparation and sowing. The strong association of feral swine with these environments is therefore expected, as these areas provide frequent access to temporary crops and residual grains remaining after harvest. Open landscapes lacking vegetation cover may also facilitate visual detection of feral swine at greater distances and are commonly monitored by local informants aware of animal activity.

Cultivated areas (CRP and DES) are typically established adjacent to riparian forest remnants within Permanent Preservation Areas (APPs; legally protected riparian buffers under Brazilian environmental law) associated with local drainage systems. This configuration forms an ideal landscape mosaic for invasive feral swine populations by combining water availability, shelter, and abundant food resources (*e.g.*, Choquenot *et al*. 1996, Herrero *et al*. 2006, Amici *et al*. 2012, Doutel-Ribas *et al*. 2019). Accordingly, water-related habitats (WRH) were strongly associated with occupancy, reflecting the species’ high dependence on water for establishment and persistence (Choquenot *et al*. 1996). Detection frequency was also positively associated with wet shrub–tree vegetation (WST), likely because these partially shaded, water-associated habitats are attractive to feral swine while simultaneously facilitating visual detection relative to dense forest habitats (Keuroghlian *et al*. 2009, Mitchell *et al*. 2009).

In contrast, wet herbaceous vegetation (WHV) negatively influenced detection frequency, likely because it represents one of the least extensive landscape classes in the state (0.4%) and provides comparatively low habitat suitability for feral swine (*e.g.*, Cordeiro *et al*. 2018). These habitats are also less accessible to local observers and informants participating in the survey.

Occupancy-based estimates indicate that feral pig populations in Mato Grosso do Sul are already extensive, with approximately 29,600 individuals distributed across invaded areas. However, density-based projections suggest that populations could potentially exceed 280,000 individuals under favorable conditions, indicating that the invasion likely remains below ecological carrying capacity in much of the state. Population reservoirs are concentrated mainly in highly productive agricultural regions (plateau) and portions of the Pantanal floodplain.

The substantial PM population estimates for the Pantanal add a critical dimension to the demographic landscape of *Sus scrofa* in Mato Grosso do Sul. Under conservative (local density) scenarios, PM populations in the Southern Pantanal (∼29,300 individuals) were comparable to the entire statewide JP population (∼29,600 individuals), while regional density-based extrapolations suggested PM populations could reach nearly 300,000 individuals—exceeding JP projections under equivalent scenarios. These estimates are consistent with historical aerial survey data for Pantanal feral pigs reported by Mourão *et al*. (2002), which documented substantial PM populations across the floodplain, providing independent validation for the demographic magnitude indicated by our extrapolation models. Taken together, these findings partially support the saturation hypothesis for long-established PM populations in the Pantanal, suggesting that this lineage has long occupied the floodplain at considerable densities, likely approaching or exceeding carrying capacity in preferred habitats.

This demographic asymmetry has important implications for the ongoing convergence between JP and PM lineages in the Pantanal. As JP populations expand from plateau regions into the floodplain, they encounter a large, long-established PM population that may serve both as a genetic reservoir and as a demographic buffer against displacement (Desbiez *et al*. 2011, Fischer *et al*. 2017, Pedrosa *et al*. 2021), while also functioning as an important bushmeat resource for local communities. The scale of the PM population also implies that hybridization between lineages, should it occur, could rapidly propagate introgressed wild boar alleles throughout a numerically dominant PM breeding pool, with potentially far-reaching consequences for local adaptation, disease dynamics, and ecosystem interactions (Mallet 2005, Herrera *et al*. 2008, Sicuro *et al*. 2025). These dynamics simultaneously highlight the demographic potential for genetic admixture in newly formed contact zones, underscoring the urgency of genetic monitoring in Pantanal transition areas where the two lineages are now converging.

The strong association between invasion intensity and agricultural landscapes suggests that continued agribusiness expansion may further facilitate demographic growth. Feral pigs responded positively to croplands, freshly tilled soils, moist shrub-tree habitats, and drainage networks, all of which provide food, shelter, and dispersal corridors. Similar relationships between resource availability and feral pig density have been documented elsewhere (Choquenot *et al*. 1996, Melis *et al*. 2006, Amici *et al*. 2012).

### Interactions between invasive feral pigs and native suiforms

Detection patterns among invasive feral pigs and native tayassuids revealed asymmetric relationships across Mato Grosso do Sul. White-lipped peccaries (*Tayassu pecari*) were consistently more frequent than collared peccaries (*Pecari tajacu*), whereas collared peccaries showed the highest number of local absences. Particularly noteworthy was the fact that collared peccaries were never detected in the absence of white-lipped peccaries in areas occupied by invasive pigs, while white-lipped peccaries persisted even under high feral pig detection frequencies. These patterns suggest greater sensitivity of collared peccaries to disturbance, habitat alteration, or invasion pressure (Cordeiro *et al*. 2018).

Although occupancy analyses detected no direct statistical relationship between native tayassuids and invasive pig occurrence, consistent with earlier findings by Oliveira-Santos *et al*. (2011) in the Pantanal, field observations indicate emerging ecological interactions. White-lipped peccaries (*Tayassu pecari*) and invasive pigs (*Sus scrofa*) frequently used the same habitats, particularly crop fields and riparian forests, although often at different times. Camera-trap records also documented aggressive encounters in which invasive pigs consistently displaced white-lipped peccaries (Laurino Pacífico, pers. comm.). Comparable interactions involving collared peccaries were not reported.

These observations suggest that invasive pigs may exert ecological effects through behavioral interference, habitat exclusion, and competition for food resources, particularly within fragmented agricultural landscapes (Gabor and Hellgren 2000, Salvador 2012, Galetti *et al*. 2015, Doutel-Ribas *et al*. 2019). At the same time, coexistence between invasive pigs and white-lipped peccaries may persist under conditions of abundant food availability, especially within highly productive agroecosystems where seasonal crops generate large resource pulses (Gabor and Hellgren 2000, Desbiez and Keuroghlian 2009, Galetti *et al*. 2015, Sicuro *et al*. 2025).

Beyond direct competitive interactions, the growing abundance and rapid expansion of invasive feral pig populations may also generate indirect sanitary consequences of significant agro-pastoral concern. Feral pigs serve as potential reservoirs for production-limiting diseases — including classical swine fever (CSF), foot-and-mouth disease, brucellosis, tuberculosis, and Aujeszky’s disease — all of which carry severe trade restrictions and economic consequences for the Brazilian livestock industry (Meng *et al*. 2009, Maciel *et al*. 2018, Kmetiuk *et al*. 2023). The risk of CSF introduction through wild boar movement has been specifically assessed for the neighboring state of Mato Grosso, where wild boars were identified as a significant pathway for disease incursion into backyard and commercial herds (Schettino *et al*. 2021), while CSF remains endemic in parts of Brazil and poses a persistent threat to the country’s disease-free zone (Robert *et al*. 2026). Bovine tuberculosis has already been confirmed in Brazilian wild boars (Maciel *et al*. 2018), demonstrating that pathogen exchange between invasive and domestic suids is not merely hypothetical. Additionally, these populations provide an abundant food source for vampire bats (Galetti *et al*. 2016), facilitating rabies virus exposure (Grotta-Neto *et al*. 2021, Teider-Junior *et al*. 2021), while native peccaries — already documented to share hemoparasites with feral pigs (Herrera *et al*. 2008) — may also be susceptible to swine pathogens due to their phylogenetic proximity to *Sus scrofa* (Kmetiuk *et al*. 2023).

## Conclusions

The invasion of *Sus scrofa* in Mato Grosso do Sul represents a fully established, large-scale anthropogenic process driven by captive breeding, agricultural expansion, and landscape connectivity. Currently occupying approximately 3.2 million hectares, invasive populations are concentrated within agricultural and hydrological corridors that provide abundant food, water, and shelter. However, the invasion remains demographically underestimated; populations are still below ecological carrying capacity across much of the state, indicating a high potential for continued demographic growth.

This expansion is already proving to be ecologically disruptive. Detection patterns and field observations indicate that native tayassuids—particularly collared peccaries (*Pecari tajacu*) — are being negatively affected through competition, behavioral interference, and habitat alteration.

Additionally, the convergence of two demographically substantial *Sus scrofa* populations—the expanding JP lineage from agricultural plateaus and the large, long-established PM population in the Pantanal— creates a critical contact zone where hybridization, competition, and transmission of production-limiting diseases such as classical swine fever and brucellosis may be amplified at a scale unmatched elsewhere in the state (Meng *et al*. 2009, Maciel *et al*. 2018, Kmetiuk *et al*. 2023). Population estimates indicate that PM populations in the Pantanal are numerically comparable to—or may exceed—statewide JP populations under equivalent scenarios, implying that the Pantanal functions as a major demographic reservoir of *Sus scrofa* with the potential to facilitate rapid introgression of wild boar alleles into the PM breeding pool. Management strategies must therefore account not only for JP control on plateaus but also for the Pantanal PM population, whose numerical dominance could both buffer against eradication and complicate genetic monitoring of admixture. Integrated genetic surveillance of contact zones, particularly in Pantanal transition areas, should be prioritized to detect and manage hybridization before admixture becomes unmanageable (Fischer *et al*. 2017, Pedrosa *et al*. 2021, Sicuro *et al*. 2025).

From a management perspective, current control efforts are critically insufficient to prevent continued expansion. Although historical benchmarks have frequently cited a universal requirement of 70% annual removal to stabilize populations (Choquenot 1998), recent critical evaluations demonstrate that effective management cannot rely on a single metric. Instead, required removal rates depend heavily on local population growth, density, and the spatial and temporal allocation of control efforts (Pepin *et al*. 2023), with populations demonstrating rapid demographic recovery following density reduction (Garabedian and Kilgo 2024). Because invasive populations in Mato Grosso do Sul remain largely below carrying capacity—a condition that facilitates high net population growth rates—control strategies must be highly adaptive, spatially prioritized, and significantly more intensive than current official rates in Brazil (IBAMA 2017, Kilgo *et al*. 2023), potentially incorporating complementary approaches such as fertility control (Pepin *et al*. 2017, Croft *et al*. 2020).

Consequently, regional eradication is no longer feasible. The persistence of this invasion reflects broader governance limitations, including weak enforcement, bureaucratic barriers, and insufficient integration among landowners, hunters, researchers, and environmental agencies, a pattern mirrored in other regions where declining hunter participation has accompanied population growth (Massei *et al*. 2015). Long-term suppression and impact reduction will require coordinated, continuous management involving stricter control of captive breeding and illegal releases, standardized monitoring programs, and integrated conservation efforts focused on protecting native fauna and vulnerable ecosystems. More broadly, the invasion of *Sus scrofa* illustrates how weak environmental governance and anthropogenic landscape transformation can convert productive agroecosystems into large-scale reservoirs for invasive megafauna, creating a continuous cycle of invasion, ecological disruption, and management failure (Figure 10).

**Figure 10.**
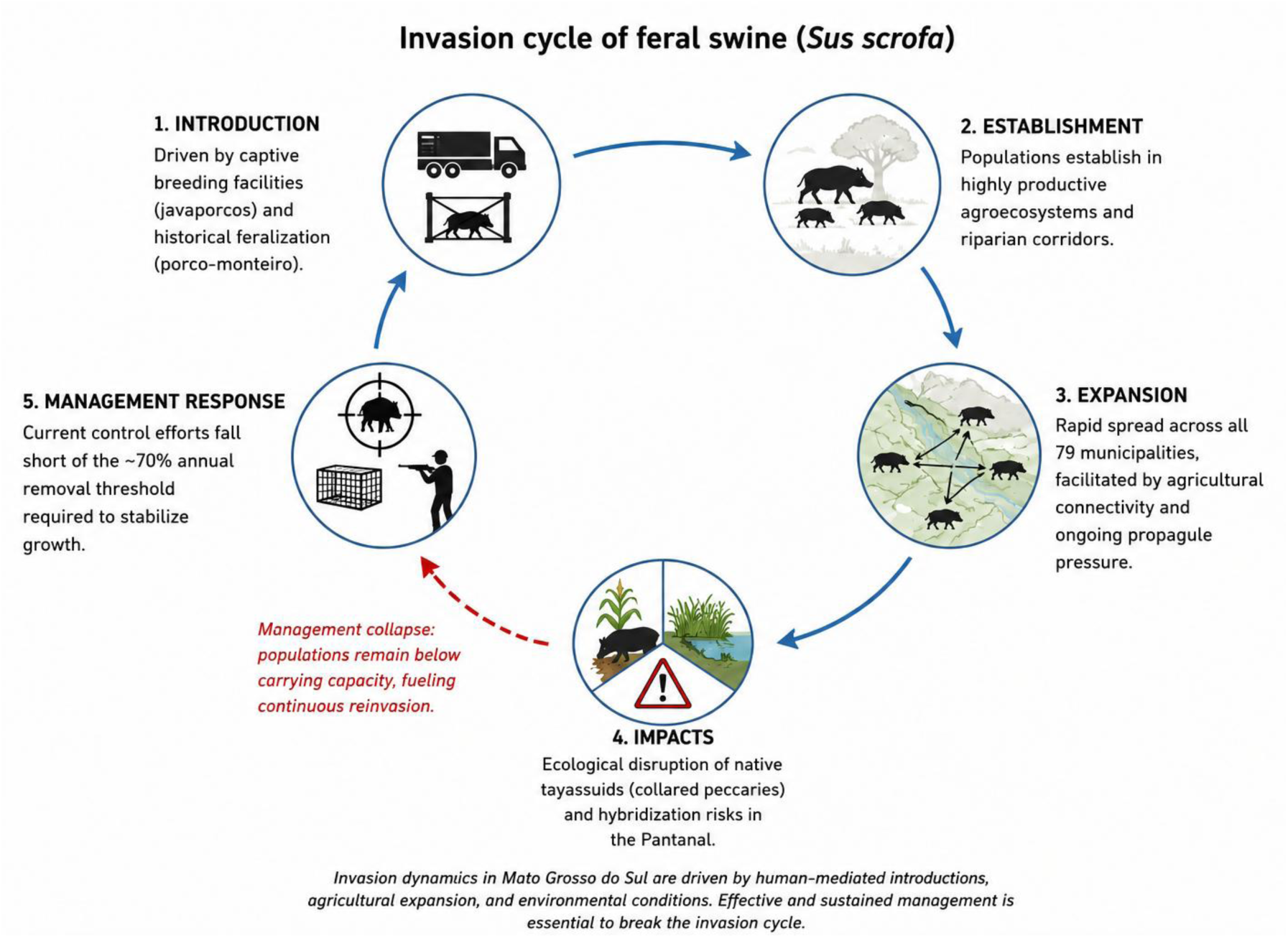
Conceptual framework of the feral swine (*Sus scrofa*) invasion cycle in Mato Grosso do Sul, Brazil, highlighting how anthropogenic drivers and insufficient management fuel continuous reinvasion and ecological disruption.

## Acknowledgments

We thank Adolfo Magariños Perez, Adriano Garcia Chiarello, Alexine Keuroghlian, Carlos Fonseca, Fábio Veríssimo Gonçalves, Geverson Dierings, José Cachimbo Nascimento, Laurino Pacífico, Luís Quinta, Michel Machado, Rafael Salerno, Rita de Cássia da Silva Paes, and Rogério Fonseca for generously sharing their expertise, field knowledge, and photographic records of invasive feral swine. We are also grateful to the regional media professionals Aline Barreto, Ariane Comineti, Evelyn Souza, Gabriela Medrado, Liana John (*in memoriam*), Marcelo Pereira, and William Franco for supporting public outreach and disseminating information related to the interviews, citizen data collection, and field investigations conducted during this study.

## Author Contributions

W.F. conceived the study, coordinated data collection, conducted statistical and ecological analyses, interpreted the results, and wrote the original and final versions of the manuscript. R.F.G.S., and A.C.P.F. contributed to geoprocessing procedures, spatial analyses, preparation of cartographic figures, interpretation of the findings, and critical revision of the manuscript.

## Use of Artificial Intelligence Tools

The authors used the Adapta ONE platform (Adapta 2026) to assist with English language refinement, data summarization, and the structural synthesis of the manuscript. Additionally, AI-assisted image editing was used to assemble and refine the comparative composite plate shown in Figure 8, based on reference photographs and under direct author supervision. This tool was employed exclusively for compositional layout and visual standardization, with no alteration of species-specific morphological traits beyond what was necessary for taxonomic accuracy. All AI-assisted outputs were carefully reviewed, edited, and approved by the authors, and all interpretations, analyses, results, and conclusions presented in this manuscript remain the sole responsibility of the authors.

## Data Availability

This study integrates historical occurrence records, interviews, institutional reports, geospatial analyses, and compiled datasets used to reconstruct the spatiotemporal invasion dynamics of feral swine (Sus scrofa) in Mato Grosso do Sul, Brazil. The datasets generated and analyzed include: (1) municipality-level records of feral swine occurrence and invasion chronology; (2) spatial layers of occupied areas and detection frequencies; (3) records of captive breeding facilities and estimated reproductive stocks; and (4) landscape and land-use variables used in statistical analyses.

Due to the sensitive nature of some information—particularly the geographic locations of informal or clandestine breeding facilities, hunting activities, and precise occurrence records associated with invasive wildlife management—raw georeferenced data are not publicly available to avoid misuse and to comply with ethical and legal considerations involving environmental enforcement and private landowners. Restricted data are available from the corresponding author upon reasonable request; requests should include a brief description of the intended use.

Non-sensitive aggregated datasets—including municipality-level occurrence records, detection frequency data, occupancy estimates, population projections, landscape and land-use variables, and statistical model outputs—will be deposited in the Dryad Digital Repository (https://datadryad.org) upon acceptance of the manuscript. A DOI will be assigned and referenced in the published article.

All analytical procedures, occupancy estimates, and statistical methods are fully described within the manuscript and its Supporting Information to ensure transparency and reproducibility. Geospatial analyses were conducted using standard GIS procedures, and statistical analyses were performed using established ecological modeling approaches. No custom software or proprietary algorithms were developed specifically for this study.

