## Supporting information: Table S1 for "Feral Pig Invasion Dynamics in Central Brazil: Human-Driven Introduction, Landscape Drivers, and Ecological Consequences"

***[Ecological Applications]***

**Table S1.** Sampling effort and respondent distribution across the 79 municipalities of Mato Grosso do Sul, Brazil, including population size, number of informants, cumulative annual sampling effort, and municipal area.

| Municipality | Population (n) | Informants (n) | Sampling effort (h·year$^{-1}$) | Area (ha) |
|---|---|---|---|---|
| **Corumbá*** | 109,899 | 36 | 11,520 | 6,496,285 |
| **Porto Murtinho*** | 16,879 | 24 | 10,240 | 1,774,441 |
| **Ribas do Rio Pardo** | 23,881 | 15 | 8,288 | 1,730,881 |
| **Aquidauana*** | 47,482 | 41 | 14,256 | 1,697,071 |
| **Três Lagoas** | 117,477 | 24 | 8,032 | 1,020,695 |
| **Rio Verde de Mato Grosso*** | 19,569 | 12 | 5,568 | 815,452 |
| **Campo Grande** | 874,210 | 42 | 16,144 | 809,295 |
| **Água Clara** | 14,992 | 20 | 12,176 | 780,921 |
| **Coxim*** | 33,323 | 14 | 6,816 | 640,922 |
| **Camapuã** | 13,694 | 22 | 7,440 | 622,962 |
| **Santa Rita do Pardo** | 7,732 | 6 | 3,200 | 613,973 |
| **Brasilândia** | 11,864 | 6 | 3,152 | 580,722 |
| **Inocência** | 7,618 | 5 | 1,952 | 577,603 |
| **Miranda*** | 27,525 | 27 | 13,856 | 547,537 |
| **Paranaíba** | 41,755 | 14 | 4,592 | 540,265 |
| **Ponta Porã** | 89,592 | 47 | 16,016 | 533,045 |
| **Maracaju** | 44,994 | 75 | 24,288 | 529,918 |
| **Sidrolândia** | 54,575 | 35 | 12,256 | 528,641 |
| **Paraíso das Águas** | 5,350 | 13 | 5,424 | 503,247 |
| **Bonito** | 21,483 | 45 | 15,552 | 493,441 |
| **Bela Vista** | 24,331 | 33 | 14,784 | 489,260 |
| **Figueirão** | 3,027 | 10 | 4,176 | 488,287 |
| **Nova Andradina** | 52,625 | 25 | 11,584 | 477,600 |
| **Alcinópolis** | 5,188 | 10 | 4,688 | 439,968 |
| **Amambai** | 38,465 | 21 | 10,048 | 420,232 |
| **Costa Rica** | 20,159 | 11 | 4,592 | 416,412 |
| **Dourados** | 218,069 | 65 | 18,608 | 408,624 |
| **Sonora*** | 18,393 | 10 | 5,056 | 407,542 |
| **Nova Alvorada do Sul** | 20,772 | 32 | 15,296 | 401,932 |
| **Rio Brilhante** | 36,144 | 50 | 17,840 | 398,740 |
| **Nioaque** | 14,092 | 17 | 8,752 | 392,379 |
| **São Gabriel do Oeste** | 25,898 | 39 | 13,808 | 386,469 |
| **Pedro Gomes** | 7,683 | 7 | 3,616 | 365,118 |
| **Cassilândia** | 21,748 | 8 | 2,464 | 364,973 |
| **Anaurilândia** | 8,927 | 11 | 4,864 | 339,544 |

| | | | | |
|---|---|---|---|---|
| **Selvíria** | 10,790 | 13 | 3,536 | 325,833 |
| **Chapadão do Sul** | 23,940 | 18 | 8,256 | 324,812 |
| **Naviraí** | 53,188 | 19 | 8,800 | 319,355 |
| **Bandeirantes** | 6,795 | 23 | 6,656 | 311,568 |
| **Iguatemi** | 15,838 | 15 | 6,576 | 294,652 |
| **Anastácio** | 24,954 | 17 | 8,336 | 294,632 |
| **Caracol** | 5,972 | 12 | 4,736 | 294,025 |
| **Jaraguari** | 7,019 | 9 | 5,232 | 291,282 |
| **Terenos** | 20,855 | 18 | 7,040 | 284,169 |
| **Aparecida do Taboado** | 25,072 | 12 | 6,992 | 275,015 |
| **Corguinho** | 5,730 | 10 | 4,496 | 263,817 |
| **Bodoquena** | 7,820 | 24 | 8,544 | 250,732 |
| **Bataguassu** | 22,389 | 17 | 6,304 | 241,760 |
| **Dois Irmãos do Buriti** | 11,132 | 17 | 6,544 | 234,165 |
| **Jardim** | 25,758 | 17 | 6,496 | 220,152 |
| **Caarapó** | 29,292 | 30 | 11,696 | 208,960 |
| **Itaquiraí** | 20,637 | 7 | 3,104 | 206,404 |
| **Ivinhema** | 23,021 | 8 | 2,464 | 201,017 |
| **Jateí** | 4,025 | 5 | 2,784 | 192,795 |
| **Batayporã** | 11,248 | 11 | 5,232 | 182,802 |
| **Rio Negro** | 4,834 | 9 | 4,032 | 180,767 |
| **Tacuru** | 11,284 | 12 | 5,648 | 178,532 |
| **Laguna Caarapã** | 7,177 | 23 | 11,120 | 173,407 |
| **Aral Moreira** | 11,771 | 9 | 2,928 | 165,566 |
| **Juti** | 6,553 | 7 | 3,104 | 158,453 |
| **Rochedo** | 5,346 | 12 | 6,016 | 156,106 |
| **Itaporã** | 23,539 | 22 | 9,264 | 132,181 |
| **Paranhos** | 13,852 | 8 | 3,520 | 130,916 |
| **Angélica** | 10,458 | 14 | 5,200 | 127,327 |
| **Guia Lopes da Laguna** | 9,991 | 12 | 6,672 | 121,061 |
| **Antônio João** | 8,808 | 13 | 6,256 | 114,518 |
| **Taquarussu** | 3,570 | 5 | 2,272 | 104,112 |
| **Coronel Sapucaia** | 15,016 | 12 | 5,184 | 102,505 |
| **Eldorado** | 12,224 | 8 | 3,520 | 101,779 |
| **Novo Horizonte do Sul** | 4,041 | 5 | 2,784 | 84,910 |
| **Sete Quedas** | 6,482 | 10 | 5,184 | 83,373 |
| **Deodápolis** | 12,773 | 14 | 7,232 | 83,121 |
| **Glória de Dourados** | 9,960 | 10 | 5,696 | 49,175 |
| **Mundo Novo** | 18,103 | 20 | 8,144 | 47,778 |
| **Japorã** | 8,836 | 6 | 2,688 | 41,940 |
| **Ladário*** | 22,590 | 6 | 3,520 | 34,077 |
| **Fátima do Sul** | 19,181 | 10 | 5,008 | 31,516 |
| **Vicentina** | 6,041 | 6 | 2,368 | 31,016 |
| **Douradina** | 5,827 | 8 | 3,392 | 28,079 |
| **Total** | **2,713,147** | **1,435** | **585,520** | **35,714,559** |

*Note. Sampling effort represents cumulative annual interview and field-survey effort expressed as hours per year (h·year$^{-1}$). Municipalities marked with an asterisk (*) partially encompass the Pantanal floodplain.*
